# A flexible quantitative phase imaging microscope for label-free imaging of thick biological specimens using aperture masks

**DOI:** 10.1101/709121

**Authors:** Chandrabhan Seniya, Catherine E Towers, David P Towers

## Abstract

A flexible quantitative phase imaging microscope is reported that offers new capabilities in terms of phase measurement from both thin and thick biological specimens. The method utilises Zernike’s phase contrast approach for label-free imaging with a Twymann-Green based phase shifting module in the back focal plane. The interfering wave fronts are manipulated by laser cut apertures to form the scattered and non-scattered fields. The design is flexible and low-cost. It is shown that the bandwidth of the optical source can be optimised to enable larger optical path differences to be measured whilst giving essentially speckle free imaging. Phase maps of the cell membrane, nucleus and nucleolus of transparent epidermis cells of *Allium cepa* have been examined as proof of concept. Measurements from a range of glass beads confirm the optical path difference capability. The implementation of the phase shifting module is *<* 10% of the cost of that using a spatial light modulator whilst delivering equivalent phase resolution.

## 1 Introduction

Cell to cell contact triggers important cell functions, hence, the ability to reliably segment and identify the boundary of an individual cell is significant to address a biological event, for example, apoptosis. However, obtaining edge features robustly is often not possible with either bright field or qualitative phase contrast methods. The most popular methods for identification of cell and cell organelles rely on the use of labelling agents, for example, fluorescence or auto-fluorescence labels, in combination with epi-fluorescent microscopy. Fluorescent labels offer molecule specific observation of events at the cellular and sub-cellular level in four dimensions and potentially at nano-scale resolution.^1^ Unfortunately, these labels have limitations due to the size of the fluorescence molecule, modification of natural biological activity, photo-toxicity and affected structural properties of the sample. In the case of imaging endogenous molecules in-vivo, labelling may not be a feasible option, which is a fundamental motivation for favouring label-free imaging techniques.^2, 3^

Recently, quantitative phase imaging (QPI)^4^ has been reported to study cell structure and dynamics by utilising Zernike’s phase contrast microscopy (PCM) approach.^5^ The quantification of cell function in a non-invasive and non-contact manner is possible at the nano-scale by measuring optical path length changes induced by cells or organelles.^6^ Common-path interferometry (CPI) configurations have been demonstrated on red blood cells,^7^ HeLa cells^8^ and for refractive index^9, 10^ and cell dry mass measurements.^11^ CPI offers superior stability for phase measurements over long time periods^12, 13^ and statistical parameters can be extracted from tissue samples.^14^ A CPI based diffraction phase microscopy method was reported^13^ but speckles generated by high temporal coherence illumination limited the phase resolution that could be obtained.^15^ Spatial light interference microscopy (SLIM) removed this limitation by employing broadband illumination and a high resolution spatial light modulator;^16^ where the spatial resolution limited by diffraction and aberrations.^17^ To study the dynamic behaviour from live cells requires a high frame rate camera and the measurements at 12.5 frames per second have been reported.^18^ The low noise QPI techniques reviewed above significantly increase the cost of the microscope owing to the use of high value components such as a spatial light modulator (SLM).

In this paper, a practical flexible quantitative phase imaging microscope (FQPIM) is described in which coded apertures are utilised to separate the scattered and non-scattered light fields. The laser cutter offers a direct advantage in that the apertures can be designed in CAD and manufactured in a range of materials with *<* 10 *μm* spatial resolution and at low cost. Therefore, FQPIM reported in this paper is very economical in the price. The approach enables any phase stepping algorithm to be implemented offering different compromises between temporal bandwidth and phase resolution performance. The source bandwidth is selected in order to give essentially speckle free imaging whilst enabling larger optical path differences to be measured than have hitherto been reported.

## 2 Principle of Flexible Quantitative Phase Imaging Microscope

The fundamental operating principle of the microscope is to utilise laser-cut aperture (LCA) masks to separate the scattered and non-scattered optical fields in the spatial frequency domain and thereby enable phase modulation. The aperture masks were produced at high resolution in order to minimise cross-talk between the two optical fields. In the case that the image fields are obtained via a Twymann-Green type interferometer two separate beams are available such that individual aperture masks can be independently optimised to produce the required scattered and non-scattered fields. This results in a non-common-path configuration and hence, the setup is sensitive to mechanical and thermal variations. These effects can be minimised by reducing the length of the arms in the interferometer and with appropriate shielding around the interferometer components, for example, beam splitter, mirror *M*_2_ and *M*_3_ (Fig. 1). An alternative is to utilize optical components where the working apertures directly correspond to the required optical fields in the spatial frequency domain; for example, a circular mirror and an annular mirror.^19^ In this case, the interferometer is common-path, however, relative movement is needed for phase modulation and hence, there has to be a greater separation (radially) between the components than for the the Twymann-Green case. Hence, it can be seen that the different setups offer different compromises; this will also effect the generation of halos^20^ where the Twymann-Green setup is expected to be superior because of the independent control over two physically separate aperture masks.

**Fig 1.**
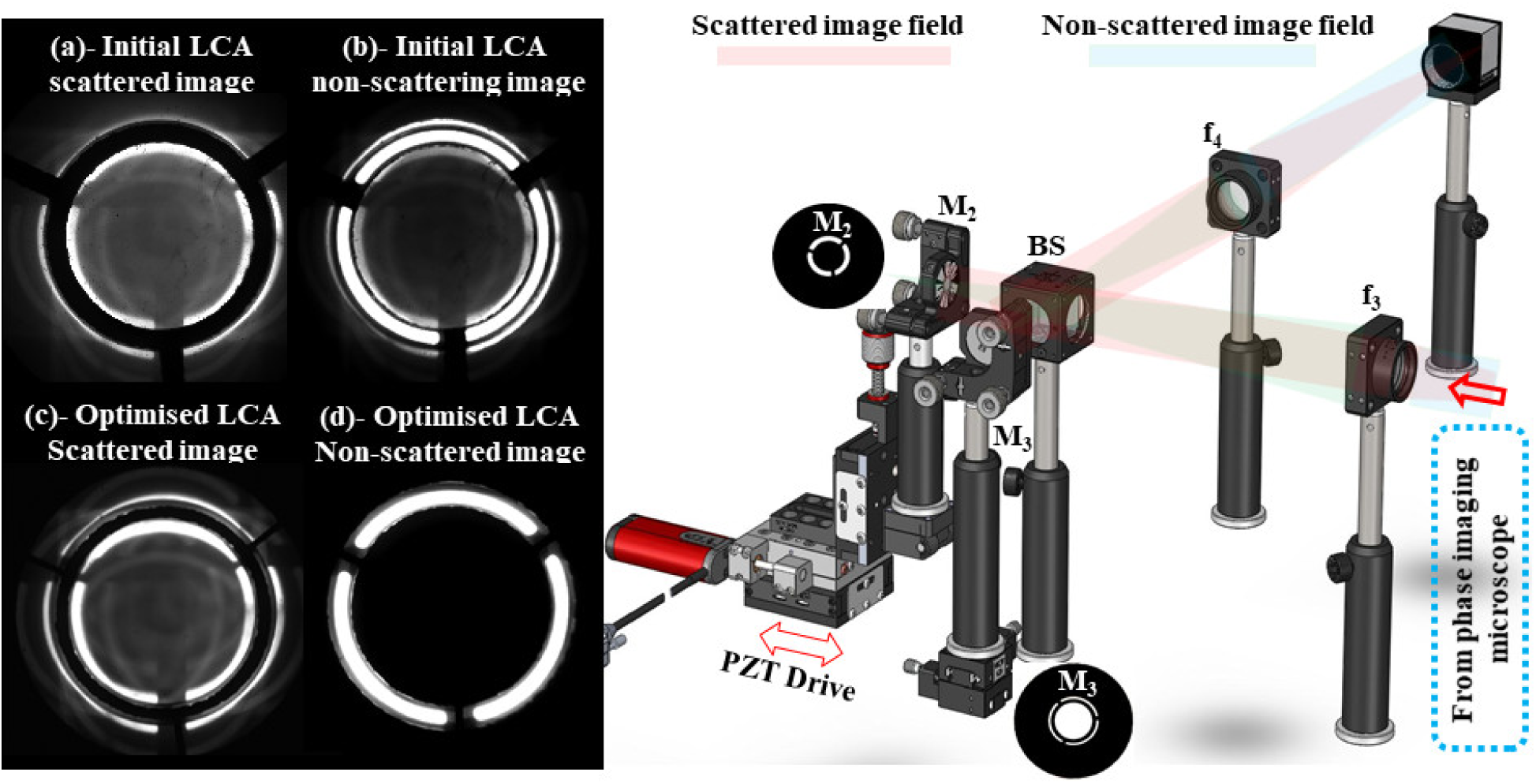
Optical Phase Shift Module in a FQPIM employing LCAs (right hand side); (a)-(b) the spatial frequency domains from the initial laser cut apertures corresponding to the scattered and non-scattered waves respectively; (c)- (d) the corresponding fields obtained from optimised apertures.

In the phase imaging microscope, the interference can be idealised as coming from the scattered and non-scattered optical fields, outputs an image plane intensity as given in Eq. 1.

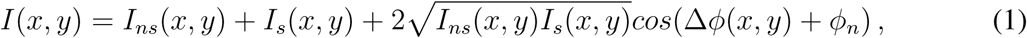

where, *I*(*x, y*) is the intensity recorded at a pixel on the CCD camera, ∆*φ*(*x, y*) is the phase difference between the scattered intensity *I*_*s*_(*x, y*) and non-scattered intensity *I*_*ns*_(*x, y*) light fields and *φ*_*n*_ is the user defined phase shift introduced between the two fields. The common-path or Twymann-Green based setups facilitate the use of a PZT mounted mirror for phase modulation; in practice, this can be considered a linear phase modulation mechanism and hence any phase shifting algorithm (PSA) can be applied directly. In this paper, the interference phase is determined from the three-frames at 120° (3F at 120°), four-frames at 90° (4F at 90°, Eq. 2) and 6+1 frames at 60° (6+1F at 60°, Eq. 3) PSAs as they offer different compromises between capture time and sensitivity to phase shifter miscalibration as defined in.^21^

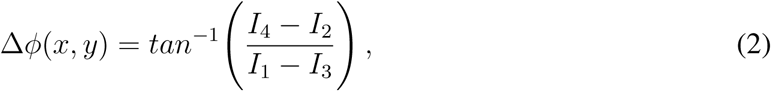

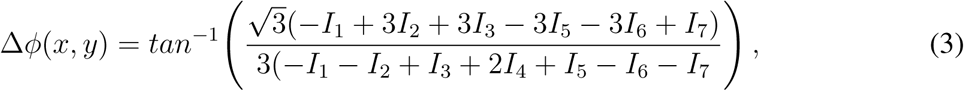

*I*_1_ to *I*_7_ are the phase shifted interference intensity images in the above equations and are the functions of (*x, y*). For each PSA, the phase is determined modulo 2*π* (wrapped) and hence, a suitable phase unwrapping algorithm^22^ is applied to determine the contiguous phase distribution under the assumption that the object phase is sufficiently sampled spatially.

## 3 Optical Configurations of LCA Mask based FQPIM

The microscope is based on an infinity corrected standard objective lens with phase contrast condenser. Köhler illumination is configured with a narrow band green LED (Thorlabs M530L3, centroid wavelength 519.90 *nm*, bandwidth 42.0 *nm* FWHM) and condenser (Olympus CX-SLC, NA - 1.25) to illuminate the sample. A non-phase contrast 10x objective (Olympus RMS10X, NA - 0.25) and the field lens (f - 180 mm, AC508-180-A-ML, Thorlabs) project the optical fields into the phase shifting module via a set of relay lenses. For the experiments reported here, the relay lenses introduce an additional magnification to give overall 25x between sample and image planes. The optical setup to spatially separate, modulate and reshape the scattered and non-scattered image fields by using laser-cut apertures (LCAs) is shown schematically at the right hand side of Fig. 1.

The complete optical design of FQPIM is presented in Fig. 2. The detailed information on size measurement and manufacturing of LCAs can be found in.^23^ The wave-fronts from the field lens are incident on lens *f*_3_ which forms part of the relay lens pair *f*_3_, *f*_4_ and hence, to form an image on the CCD camera. Between the relay lenses is the interferometer where the two mirrors *M*_2_ and *M*_3_ are placed in the spatial frequency domain and have LCAs applied to their front surfaces. A VLS6.60 laser cutter (Universal laser systems) was used in this case offering a spatial resolution of 0.03937 pixel/*μm* and the apertures were cut in matte black paper (thickness 0.20 *mm*). The aperture applied to *M*_2_ is designed to isolate the non-scattered waves corresponding to the NA of the condenser annulus. The aperture at *M*_3_ is configured such that the scattered waves at an NA lower and higher than that of the condenser annulus are reflected. The operation of each laser cut aperture was confirmed by inserting additional relay lens optics to form an image of the spatial frequency plane from each mirror in turn, see Fig. 1a) -d). The geometry of the initial LCAs were determined by calculation to give the optical fields shown in Fig. 1a) and b) for the scattered and non-scattered fields, respectively. In particular, there is cross-talk of the scattered light being observed in Fig. 1b) which is intended to reflect only the non-scattered waves. Optimised geometry laser cut apertures were identified from these images resulting in the scattered and non-scattered fields in Fig. 1c) and d), respectively. It can be seen that the cross-talk present in Fig. 1b) has been removed in Fig. 1d). The scattered light seen in Fig. 1c) is from an increased area of the spatial frequency domain than in Fig. 1a) and hence, also has greater amplitude. Interference contrast can be maximised by balancing the irradiance of the scattered and non-scattered image fields and can be achieved by further reducing the open area of the aperture for the non-scattered waves (at *M*_2_, Fig. 1d). Phase shifts between the two waves are obtained by a piezoelectric transducer (PZT, Thorlabs, PK3JMAP1, *£*43.54) installed in a high precision linear stage, thus the system is low-cost. Furthermore, the lateral resolution is determined by the objective as the NA of the scattered waves is not restricted in the phase shift module.

**Fig 2.**
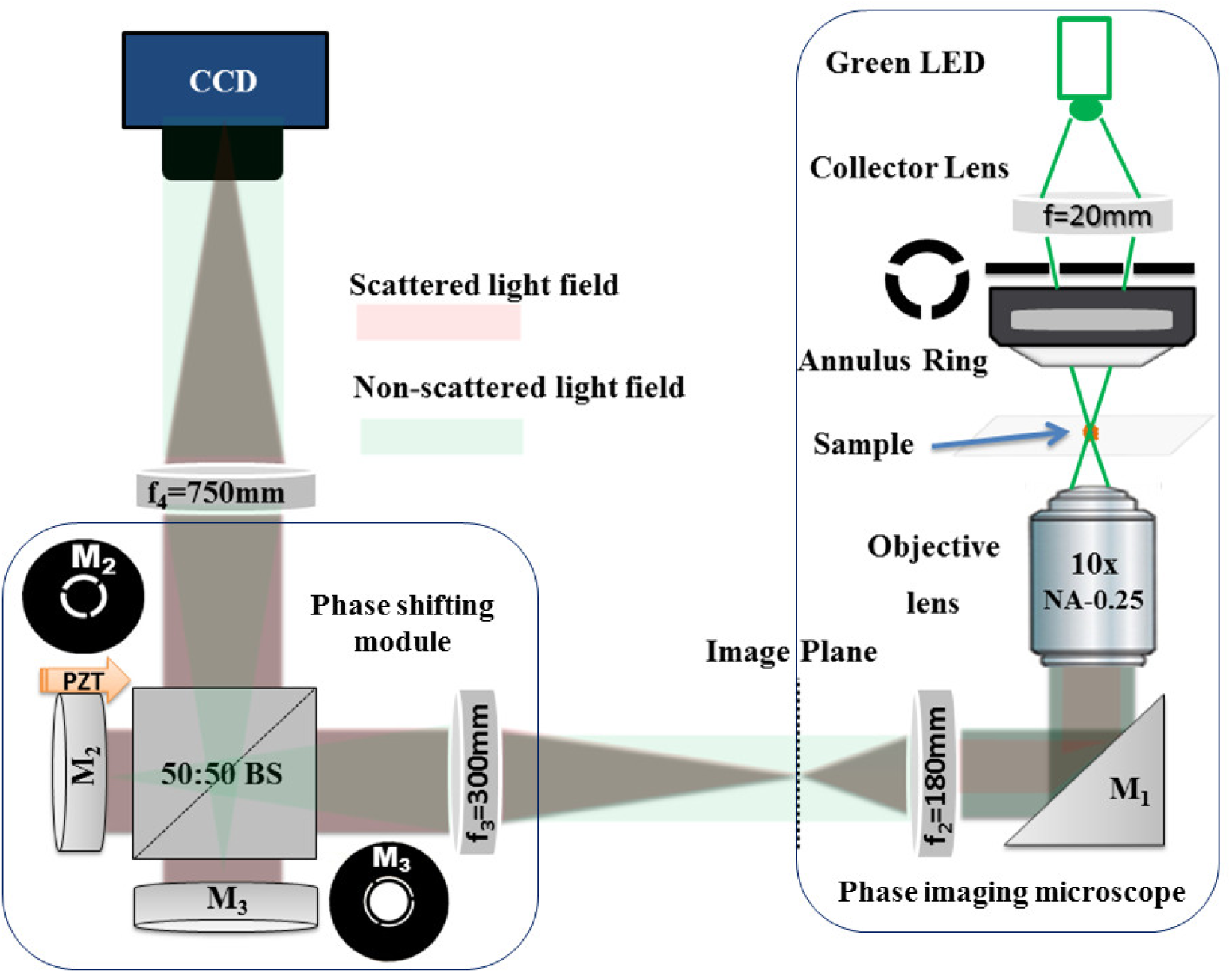
The non-common-path optics for phase shifting in FQPIM based on LCAs; *M*_1_ *− M*_3_ (mirrors where coded LCAs are applied), and *f*_3_ − *f*_4_ (relay lenses).

## 4 Source Characteristics for Broadband Phase Shifting Interferometry

Coherence theory for quasi-monochromatic light shows that the interference visibility depends on the product of spectral bandwidth and time delay, i.e. optical path difference, between the two interfering beams.^24^ Many optical sources can be approximated as a Gaussian function with respect to optical frequency, *ν*, in which case the visibility *V* (*τ*) is given by [Falaggis]:^25^

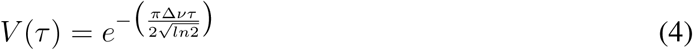

where, 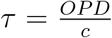 is the time delay due to the optical path difference (OPD); c is the speed of light and ∆*ν* is the FWHM frequency bandwidth of the source. It is recognized that recent trends in microscopy are towards broad bandwidth sources covering most of the visible spectrum, however, this would then limit the maximum OPD measurable due to reductions in fringe visibility. A graph is given in Figure 3 showing the maximum frequency bandwidth compatiable with measuring an OPD of 1500 nm with a fringe visibility of 1*/e* and for a range of centre wavelengths of a Gaussian source. Overlaid on this graph are results of a computational simulation for the phase analysis process applied to a 1-D increasing OPD ramp from zero (0) to the maximum value, in this case 1500 nm. The phase analysis includes generating the recorded intensities corresponding to the 4F at 90 phase stepping algorithm, with shot noise modelled as the primary noise source and 1-D phase unwrap along the increasing OPD axis.^26^ The success, or otherwise, of the phase analysis is emphasised by looking at the percentage of simulations that contain unwrap errors by comparing the unwrapped phase with the known 1-D phase ramp (the vertical axis in Fig. 3). The correspondence between the theoretical limiting source bandwidth and the simulation results for successful phase analysis can be seen in Fig. 3. For bandwidths less than the theoretical limit, the phase unwrapping process gives 0% errors, whereas for bandwidths larger than the theoretical limit the percentage of unwrap errors rapidly increases to 100%. Additional vertical lines are shown on the graph to indicate typical optical sources in microscopy. It should be noted that for a lower OPD limit, e.g. 250 nm, the phase analysis process gives 100% success over the same range of centre wavelength and FWHM bandwidth as given in Fig. 3. These results indicate that narrowband LED sources with *<* 50 nm FWHM bandwidth are suitable for large OPD measurement in phase stepping interferometry for microscopy; with greater bandwidths being applicable for longer centre wavelength sources. The position of the green LED (Thorlabs, M530L3) used in this paper is indicated.

## 5 Results

### 5.1 Validation and Phase Sensitivity Assessment of FQPIM

The results presented were obtained using the Twymann-Green based system employing LCAs as shown in Fig. 1 and Fig. 2. The performance of the microscope was evaluated by imaging a test target (R1L3S5P, Thorlabs) and the glass beads. Approximately uniform phase areas were selected for the assessment of the phase resolution where deviations from a low order polynomial fit (spatially) can be used to determine a standard deviation phase noise (following the method described in^27^). The representative unwrapped phase map is shown in Fig. 4 and microscope’s performance data from 6+1F at 60° and 4F at 90° PSAs are given in Table 1.

**Table 1.** Phase Resolution Assessment of the FQPIM.

| Assessment Parameter | 4F at 90° PSA | 6+1F at 60° PSA |
| --- | --- | --- |
| Mean phase shift angle ($\alpha$, degrees) | 90 | 60 |
| Std Dev in phase shift ($\sigma$, degrees) | 1.94 | 2.58 |
| Phase noise for column 1077 ($\sigma_\phi$, fringe fraction) | 1/1338th | 1/1664th |
| Phase resolution for column 1077 (nm, $\lambda$ - 530 nm) | 0.382 | 0.317 |

**Fig 3.**
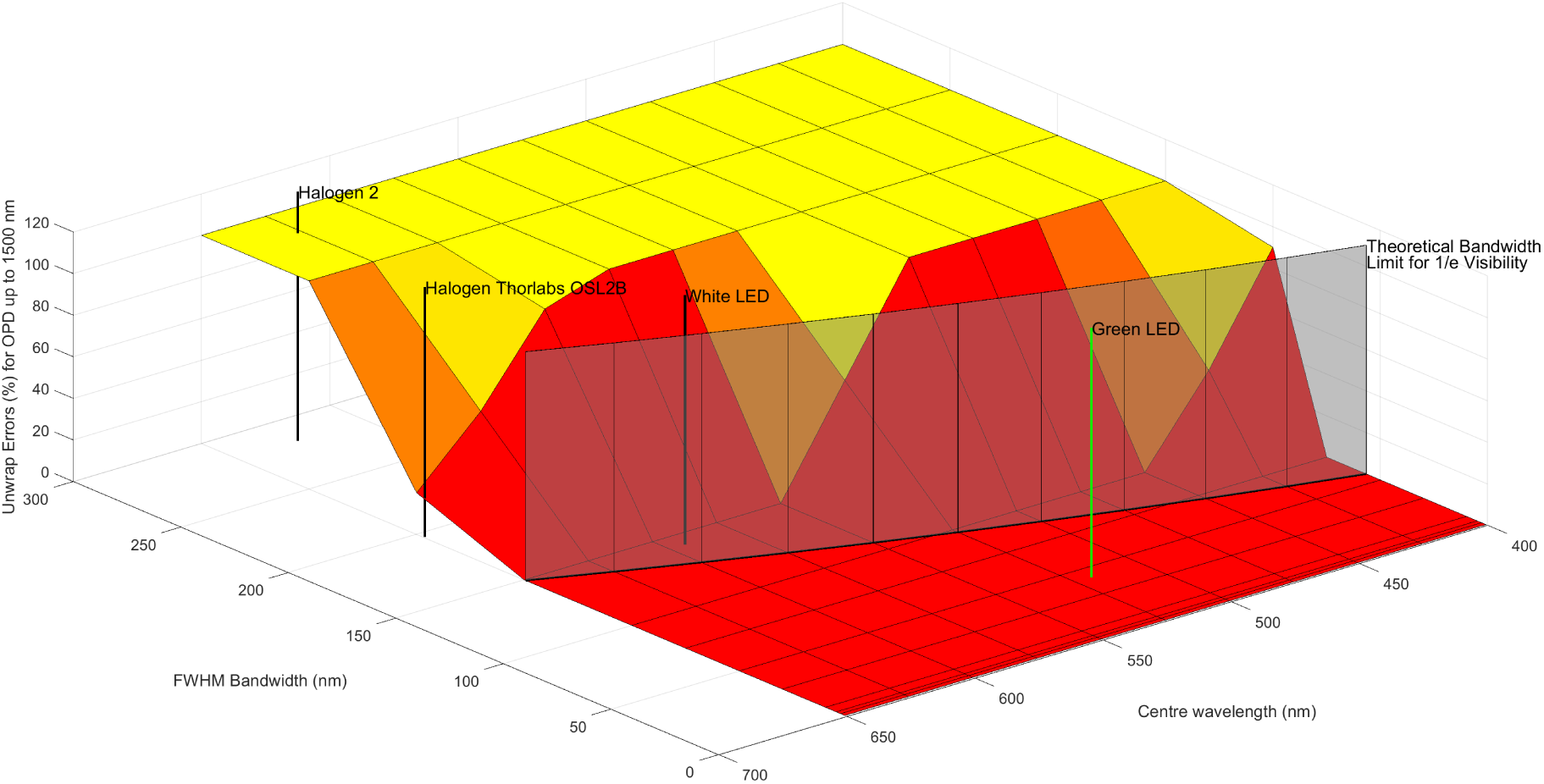
Simulation of Phase unwrapping performance up to an OPD of 1500 nm as a function of centre wavelength and FWHM bandwidth of a Gaussian intensity profile optical source. High values (yellow) indicate 100% phase unwrap errors, low values (red) indicate successful 0% unwrap errors. The theoretical bandwidth limit for interference visibility of 1/e is indicate by the semi transparent grey surface. The position of typical light sources used in microscopy are indicated.

**Fig 4.**
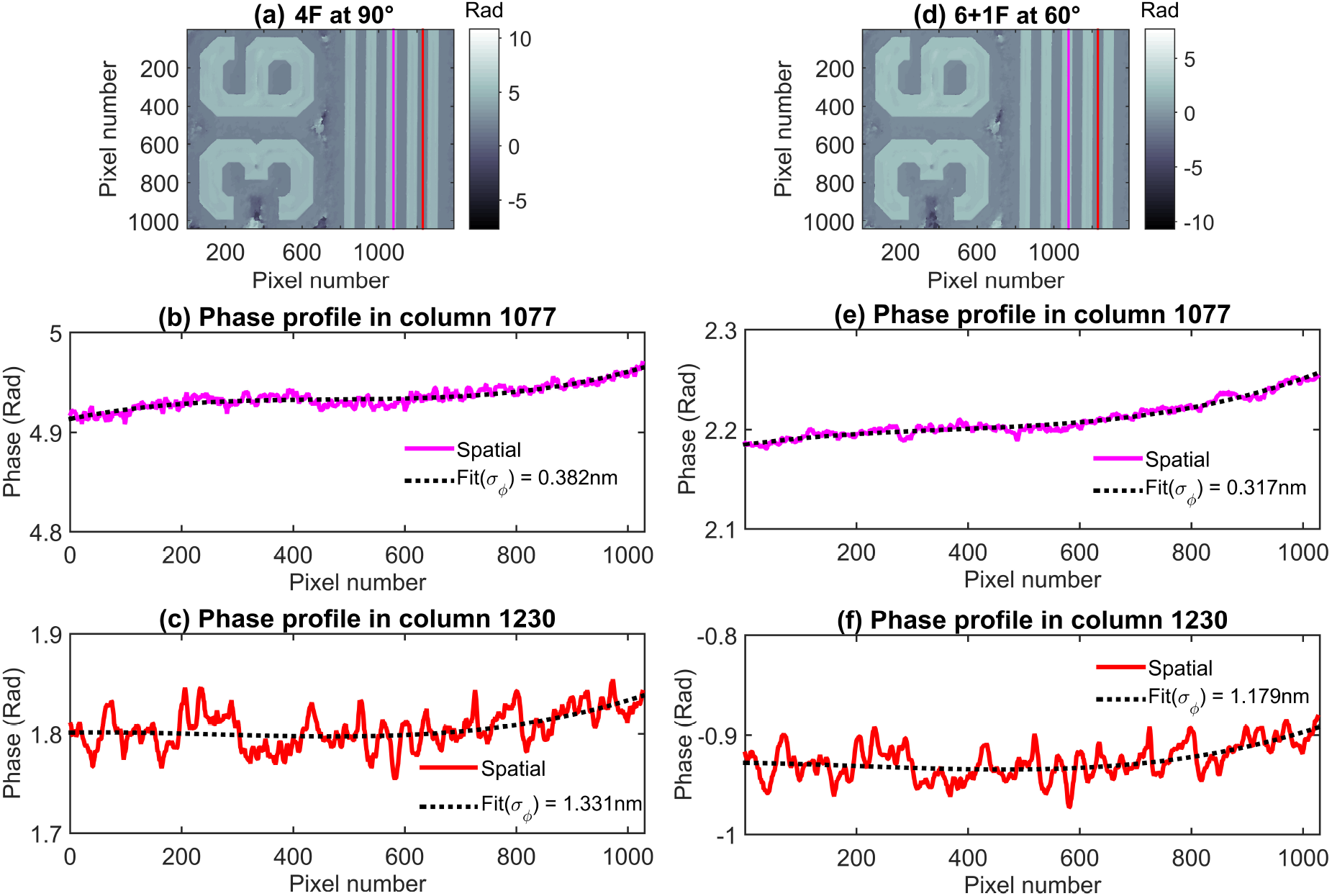
a), d) Unwrapped phase maps of test target obtained under green LED illumination with the 4F at 90° and 6+1F at 60° PSAs respectively. b), c), e), f) Phase profiles from approximately uniform regions with low order polynomial fits (black dotted curves) and hence, the standard deviation phase noise is obtained and indicated in each graph. Data from column 1077 is indicated in pink and that from column 1230 in red.

In Fig. 4(b-c and e-f), phase measured in radians was converted to *nm* using Eq. 5 for the sensitivity analysis, where, *λ* is the centroid wavelength of an illumination source and OPD is the optical path difference.

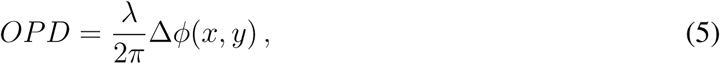

Using this method, a direct measurement of the OPD changes introduced by the test object relative to the reference field is possible. ∆*φ*(*x, y*), is directly proportional to the integration of refractive index *n* along the direction of propagation. For a glass bead and considering the ray passing through the bead centre, thickness *t*(*x, y*) of the glass bead (diameter) with refractive index *n*_*gb*_ and that of the surrounding fluid, for example glycerol *n*_*gly*_ are mathematically related as:

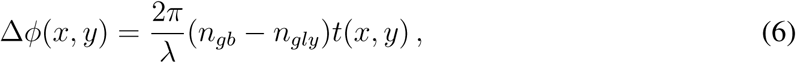

It can be seen that the PZT calibration was achieved to similar accuracy for each PSA and for both algorithms the phase resolution performance is better than 1/1300^*th*^ of a fringe. The higher number of frames utilised by 6+1F at 60° PSA yields substantially better phase resolution than for the other algorithms considered. Equally important is the modulation depth in regions that are distant from the sample and hence, where the scattered waves are weaker, for example the phase resolution in column 1230 of Fig. 4 compared to that in column 1077. The local phase resolution depends on the intensity of the two beams and noise in the input images and is expected to degrade in regions away from scattering objects.

A small pinch of glass bead powder (10 - 30 *μ*m, refractive index - 1.51, Polysciences, Inc.) was thoroughly mixed in glycerol (refractive index - 1.46572) to generate a sample containing a broad range of OPDs. The *x − y* size of individual bead images can be used to determine the diameter of each bead. The spatial scale factor in the images is known theoretically from the objective and relay lenses giving 25x magnification overall. Images were also obtained from an NBS 1963A resolution chart (Thorlabs, part number R1L3S5P) showing *<* 0.1% error in expected magnification. The phase profile of an isolated bead within the sample is given in Fig. 5 using four PSAs.

**Fig 5.**
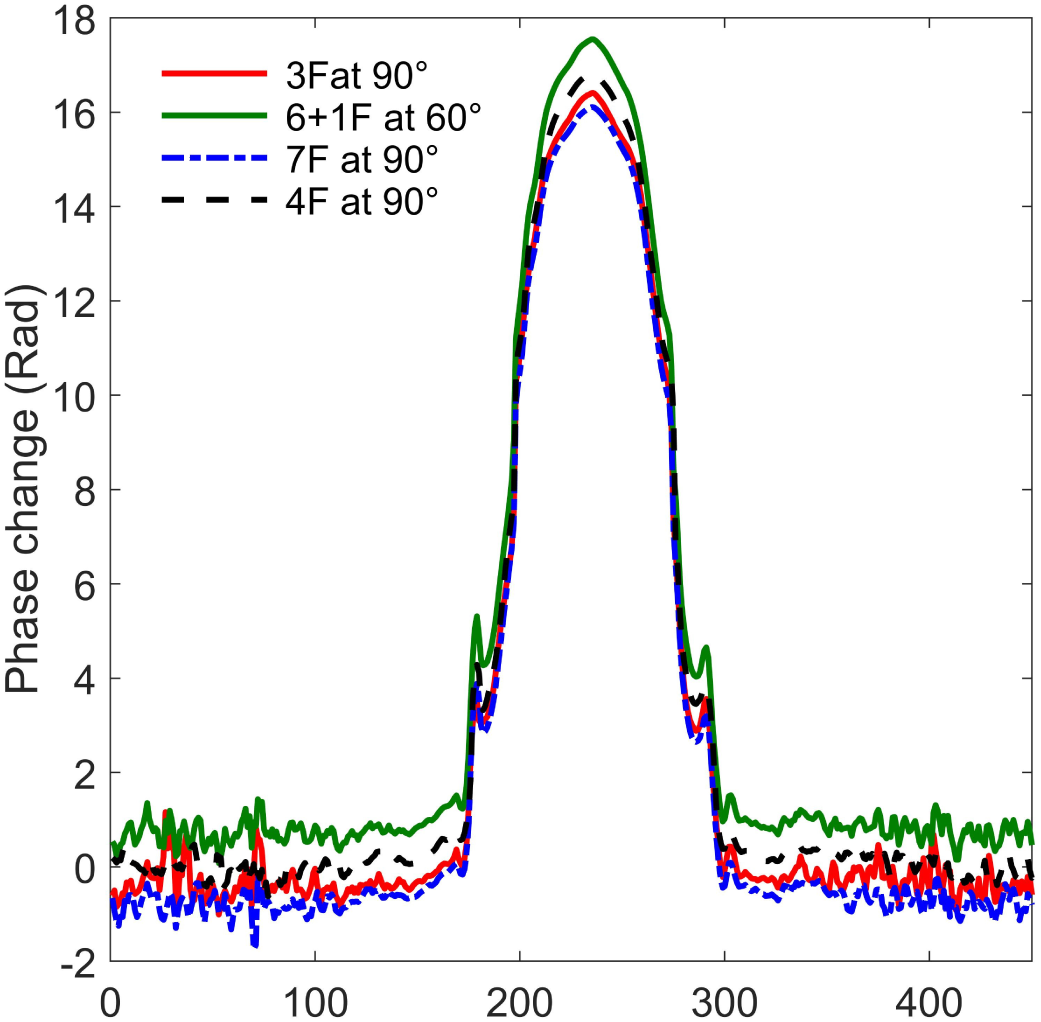
Phase profile of thick glass bead against pixel number on the CCD camera.

Phase offsets have been applied as appropriate in order to separate the results from the different PSAs. The phase difference between the background and bead centre is consistent for all PSAs at approximately 16.5 radians, i.e. 2.63 fringe orders. The phase gradient around the edges of the beads is steep and can cause bending of the light waves in this region. The steep gradients at the bead edge are a cause for concern for the phase unwrapping algorithm; however, as can be seen from the experimental phase profiles in Fig. 5, the deviations observed are not of sufficient magnitude to give an unwrapping error. Furthermore, in PCM, the halo or shade off effect is a general problem in the phase images due to overlapping of the scattered and non-scattered light fields in the Fourier plane.

A set of 4 glass beads were isolated for further analysis and their unwrapped phase difference, bead centre to background, is plotted against the measured *x − y* diameter in Fig. 6 (values: 11.93 radian, 12.17 radian, 4.97 radian and 16.54 radian were measured for 23.478 *μm*, 23.99 *μm*, 29.412 *μm* and 31.476 *μm* beads, respectively). A linear fit to these measurements (black dashed line) gives an *R*^2^ value of 0.999. Theoretical values for the phase difference were also calculated from Eq. 6 using manufacturers data for the bead and glycerol refractive index and the measured *x − y* diameters for the thickness (red line in Fig. 6).

**Fig 6.**
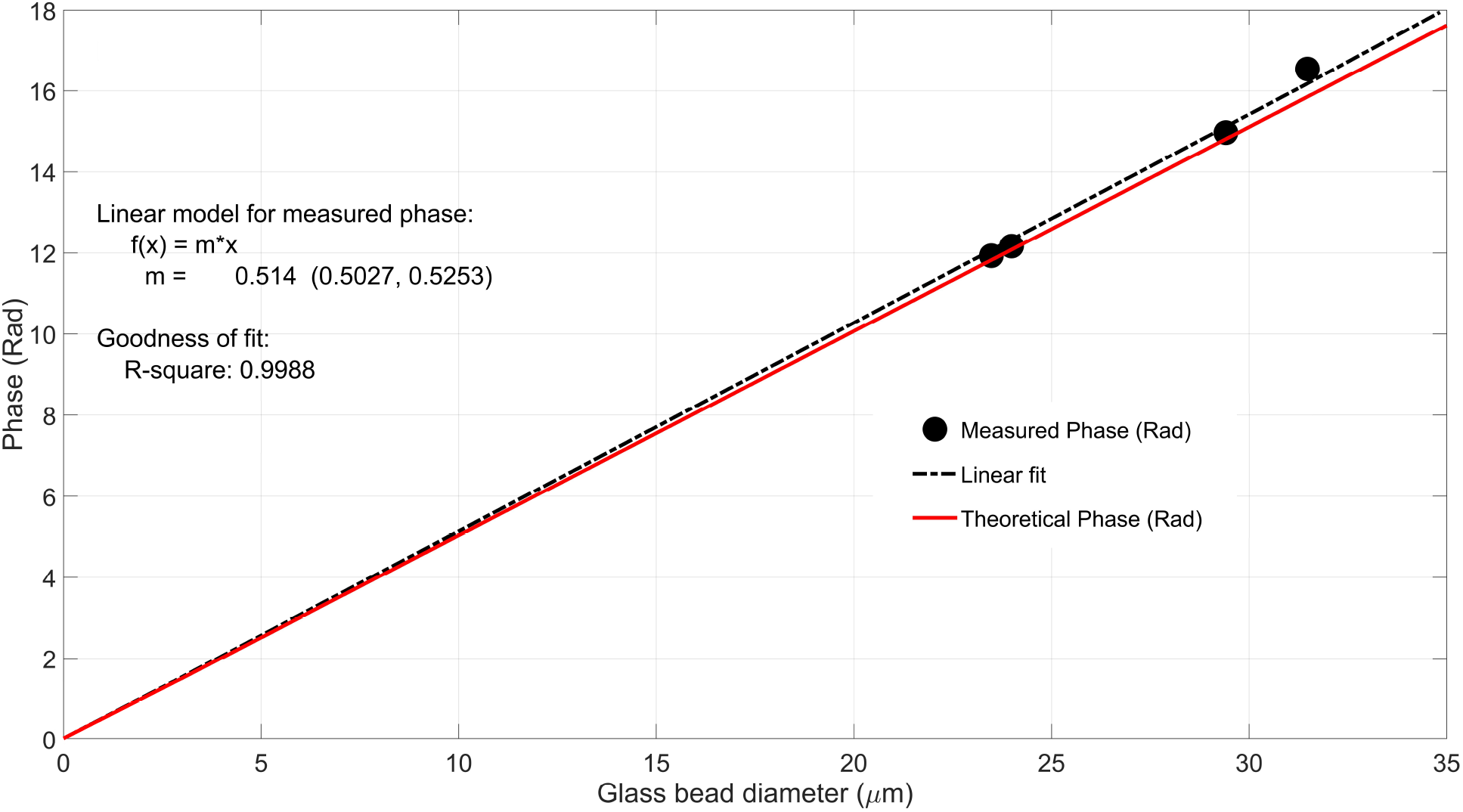
Theoretical validation of phase measured from glass beads.

An uncertainty of 2.065% was measured between the theoretical and experimental phase measurements from the best fit slope. An uncertainty analysis of Eq. 6 shows the fractional uncertainty in the unwrapped phase as:

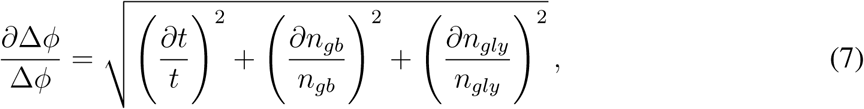

Assuming that the uncertainty in diameter measurement from the (*x, y*) image is 1 pixel at either edge and the uncertainties in the refractive indices are half the least significant digit of the values quoted then the fractional uncertainty in the unwrapped phase is estimated as 2.03%. This is approximately consistent with the uncertainty from the difference in slopes between the measured and theoretical data. The measured phase differences appear to be systematically higher than the theoritical predictions. This is believed to derive from the assumed value for the refractive index of glycerol which is strongly dependent on low levels of water concentration (5% water content gives a refractive index change of 0.008)^1^.

### 5.2 Assessment of Temporal Phase Stability of FQPIM

The temporal phase stability of the microscope has been quantified by recording 100 sets of 4F at 90° interference images of the test target (R1L3S5P, Thorlabs). A representative section of a wrapped phase map is shown in Fig. 7 (left) which identifies 4 probe points. The temporal phase evolution at each probe point is given in Fig. 7 (right) showing highly correlated variations consistent with drift in the interferometer section of the microscope. A temporal drift of 0.4 radians or approximately 1/15^*th*^ of a fringe occurred in the 3 minute data capture time. Such drift can be compensated in practical, long time series measurements by having a suitable time stationary part of the object typically at the edge of the field of view.

**Fig 7.**
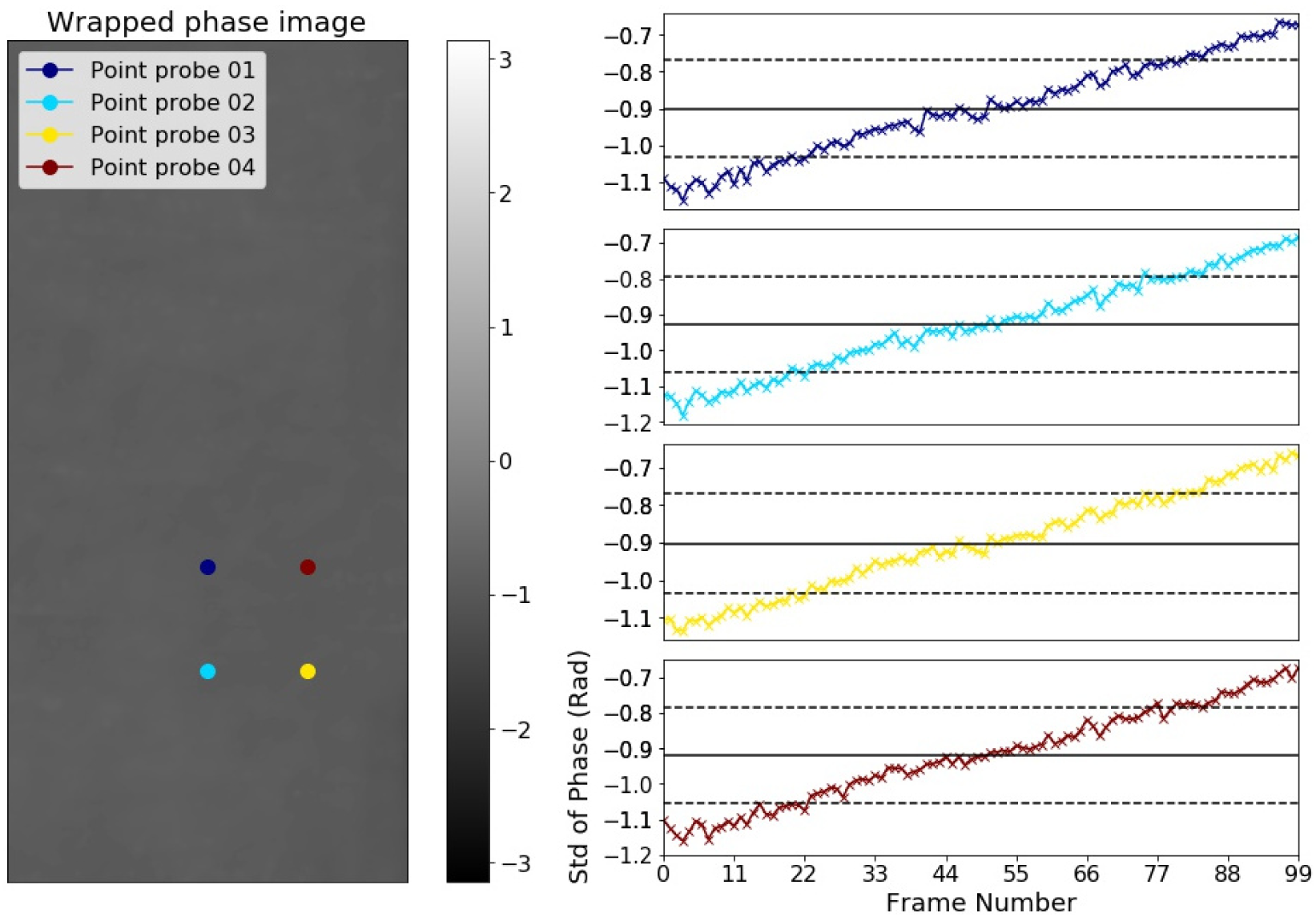
Left - Unwrapped quantitative phase image of test target (Colorbar in radians). Right - Selected four point probes for temporal phase noise over 100 wrapped quantitative phase images.

### 5.3 Phase Measurement from a Biological Sample

A slide of semi-transparent unstained epidermal cells of *Allium cepa* (Instruments Direct Services Ltd, Product code: MSAS0111) was used to assess the capability of the microscope to examine a complex biological objects. The sample is essentially a phase object having little effect on the transmitted amplitude. Fig. 8 demonstrates the microscopes ability to quantify the phase changes induced by spatially fine objects, for example, the nucleus and the nucleolus at sub-nanometer scale. The broadband illumination from the green LED provides speckle-free image light fields and hence, a good spatial resolution is achieved. The cross sectional line profile of phase for the line 1, 2 and 3 are shown in Fig. 9 and Fig. 10. The nucleus cross section and the line 1 profile in Fig. 8 also presented in Fig. 10 for 3 PSAs. It can be seen that there is good consistency between the PSAs (it should be noted that the unwrapped phase from the 3F at 120° PSA has been offset compared to the other two PSAs), despite the complexity of the cells as phase objects containing step changes at the cell wall. However, the unwrapping algorithm produces different results in particular spatial regions, for example, the box indicated to the right-hand side of profile 2 in Fig. 9a). It is interesting to note that PSAs with higher numbers of images tend to be more robust against phase unwrapping failures which may arise due to combinations of modulation between reference and object beams, and mechanical vibrations.

**Fig 8.**
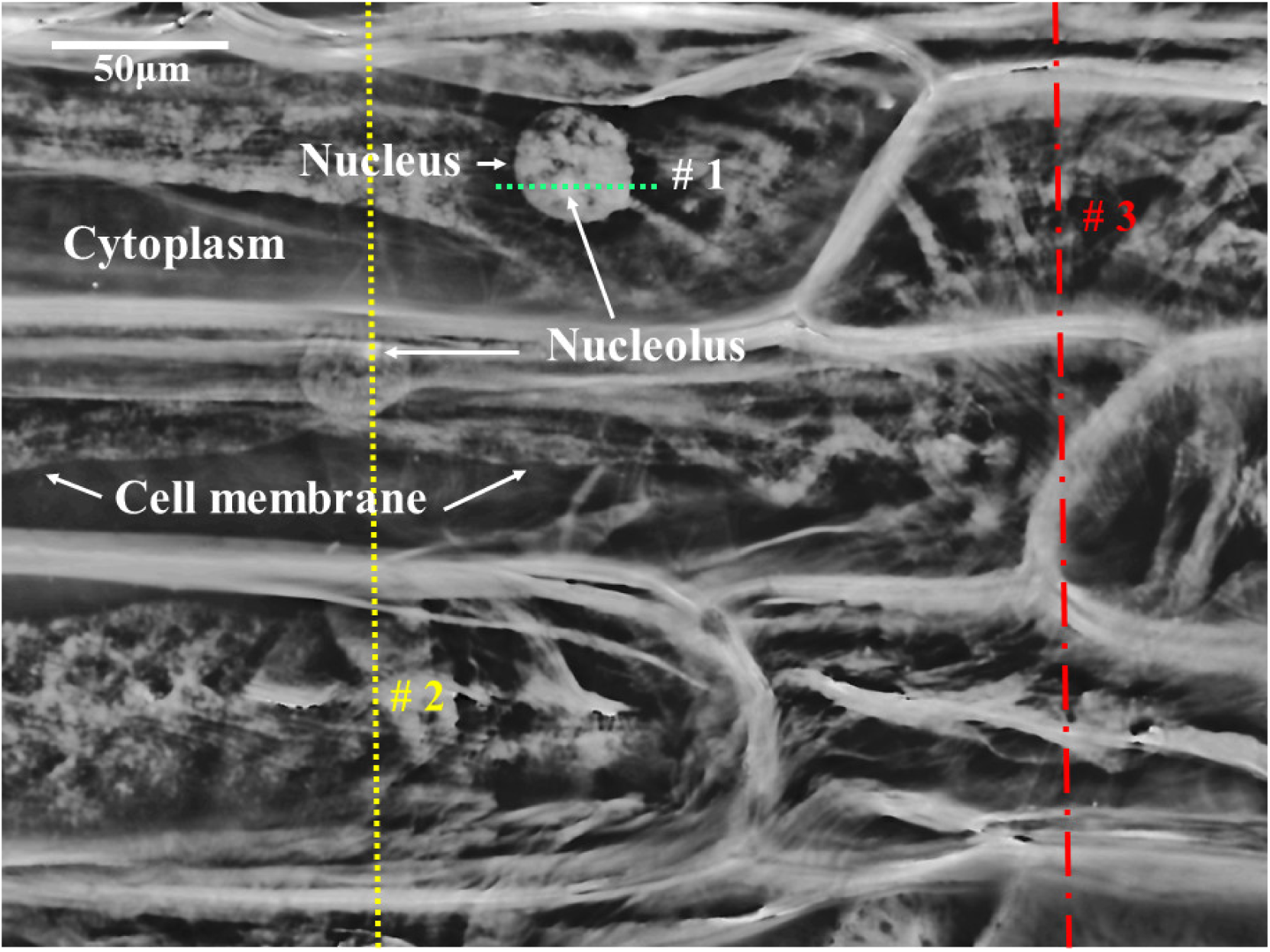
Unwrapped phase map of epidermis cells of *Allium cepa* using 6+1F at 60° PSA, line 1, 2 and 3 areas are selected for further assessment (see, Fig. 9 and Fig. 10).

**Fig 9.**
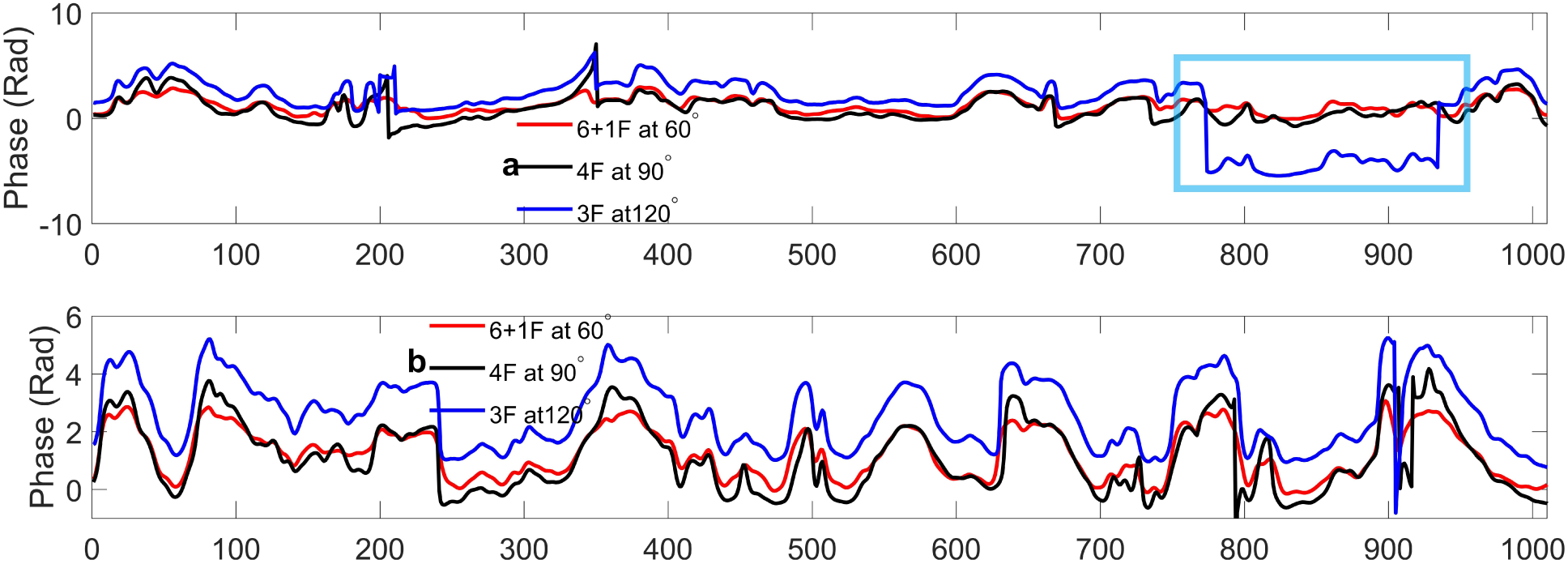
Cross sectional profile of the line 2 (top) and line 3 (bottom) mentioned in the Fig. 8.

**Fig 10.**
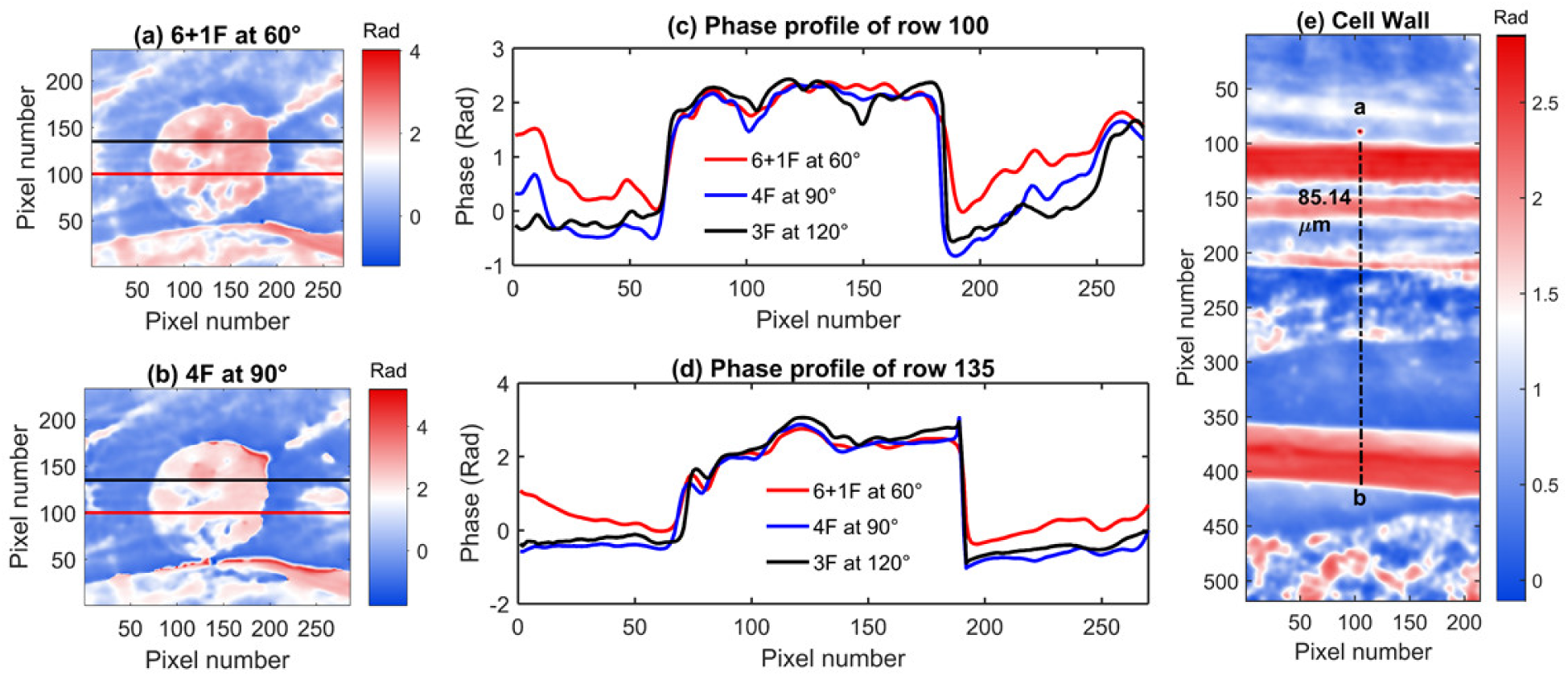
a) - b) The unwrapped phase map of the nucleus of *Allium cepa* using 6+1F at 60° and 4F at 90° PSAs, respectively. c) - d) Phase profile through the nucleus for the row 100 and 135 using 3 PSAs. e) Measured cell wall.

The phase change induced by the nucleus has been quantified as 2.343 radians (see, Fig. 10), with an additional phase corresponding to the nucleolus of 0.418 radians. These data reinforce the requirement for high phase resolution as the nucleolus corresponds to a phase feature of approximately 1/15^*th*^ of a fringe. From the lateral dimensions in the image, the nucleus is observed to be approximately 30 microns in diameter Fig. 10a) and b).

The literature indicates typical refractive indices of the cell wall (*n* = 1.41*−*1.52) for flowering plants,^28^ cytoplasm (*n* = 1.36 *−* 1.39), nucleus (*n* = 1.355 *−* 1.65) and nucleolus (*n* = 1.375 *−* 1.385) for animal cells.^29, 30^ These values are numerically consistent; however, it should be noted that the uncertainty in refractive index limits the ability to accurately quantify the thickness of organelles in many cases; additionally, preparation of the sample between microscope slides may also affect the thicknesses measured. From the lateral information the cell wall is typically 13.674 *μ*m thick and the cell has a width of 85.24 *μ*m (see, Fig. 10) which are within the expected range.

## 6 Summary

In this paper a flexible quantitative phase imaging microscope has been presented. The optical setup employs laser cut apertures to obtain the wave fronts corresponding to the scattered and non-scattered fields in the Zernike’s phase contrast arrangement. As the apertures are simple to produce and the geometry can be defined at high resolution it is possible to fine tune the cross-talk between the two waves and hence optimise microscope performance. The theoretical analysis of the light source bandwidth and measurable OPD shows that for thick biological samples narrow band LED sources are required in order to maintain interference contrast. These data have been verified by a computational model of the phase analysis process. Measurements from glass beads have verified the large, 16.5 radian OPD capability of the microscope. Data from a biological sample has been presented confirming the ability to resolve sub-cellular features with high lateral and phase resolution performance consistent with speckle free images. The cost of the phase shift module of the microscope presented here is *<* 10% of that of the widely reported SLM based quantitative phase measuring microscopes.

### Disclosures

The authors do not have any conflicts of interest.

## Acknowledgments

**Chandrabhan Seniya** thanks the Madhya Pradesh State Govt. of India for an Overseas Fellowship and the School of Engineering, University of Warwick, United Kingdom for support during his PhD. The authors acknowledge financial support from the Medical Research Council through grant MR/K015613/1.

## Footnotes

1 http://www.aciscience.org/docs/physical properties of glycerine and its solutions.pdf. Accessed on 04/10/2017

